# A simple tool extends TIMSTOF compatibility with historic data processing tools and enables ion mobility-enhanced spectral libraries

**DOI:** 10.1101/2021.10.31.466659

**Authors:** Benjamin C. Orsburn

**Affiliations:** The Department of Pharmacology and Molecular Sciences, The Johns Hopkins University School of Medicine, Baltimore, MD 21205

## Abstract

Trapped ion mobility mass spectrometry is proving to be a disruptive technology in LCMS based proteomics. One primary drawback of this hardware is the lack of compatibility with the hundreds of data processing pipelines historically in use. This study describes a simple data conversion tool that “folds” the TIMSTOF ion mobility data into the MS2 fragmentation spectra allowing simple downstream processing. Little to no detriment in the assignment of peptide spectral matches is observed when “folding” the 1/k0 value into the low mass region. To demonstrate one utility of TIMS Folding, spectral libraries are provided in multiple common formats that were constructed from the same files both with and without folded ion mobility data. When new data is acquired and folded using the same parameters prior to data processing the folded ion mobility data can be used as an additional metric for peptide match confidence against folded spectral libraries.

## Introduction

Trapped ion mobility allows the near simultaneous separation and accumulation of ionized molecules prior to time of flight mass spectrometry.(Meier et al., 2018) Coupling ion accumulation with simple and rapid quadrupole time of flight hardware allows high relative scan rates while maintaining sensitivity higher than any previously described time of flight instruments. Today only a limited number of over 1,000 proteomics data processing software packages in use today (Tsiamis et al., 2019) can process TIMSTOF data due to both the high relative data density and unique nature of the acquired data. While an increasing number of software packages now accept TIMSTOF data, compatibility may not directly correspond to ease of use, as even the fastest search tools require significantly more time to process each TIMSTOF file.(Yu et al., 2020a, 2020b) While it is possible to extract simplified text output from each MS/MS spectra by converting the vendor proprietary format to universal formats, only the historic Mascot Generic Format leads to an overall decrease in data size. Conversion of the vendor .d format to mzXML, for example may lead to a dramatic increase in the size of each file. When files are converted to MGF the ion mobility measurements are no longer accessible by most software packages.(Martens et al., 2011; Turewicz and Deutsch, 2011) During the creation of MGF text files it is possible to retain the 1/k0 value of each parent ion by incorporating that value into the scan header, as in the default PASEF MGF conversion settings in recent versions of MSConvert. To exploit this I have developed a lightweight program that embeds the 1/k0 value from the scan header into the corresponding MS/MS spectra by adding this value to a mass of the user’s choice. By placing the new fictitious ion mobility fragment ion in the low mass region that is typically ignored by data processing tools, this modification causes no significant change in the peptide spectral matches obtained.

The “TIMSFolding” process creates a new MGF that is only subtly altered from the original file and can be processed in historic programs that can utilize the MGF file format. While this is an early work that is under progress, the integration of the 1/k0 value has proven useful in providing an additional confidence metric for each peptide spectral match obtained.

## Methods

TIMS Folding was compiled in C++ in Visual Studio 2017. The source code and executable are freely available on GitHub at the following link. https://github.com/orsburn/TIMSFolding

### Generation of K562 Proteomics Data

A single injection of 200 nanograms of K562 digest from Promega was injected on an EasyNLC1200 coupled to a TIMSTOF Flex mass spectrometer linked by a 25cm IonOpticks Aurora nanoLC column. A 60 minute gradient with a flowrate of 200nL/min was used which began at 95% Buffer A (0.1% formic acid in LCMS grade water) 5% Buffer B (80% acetonitrile with 0.1% formic acid). Both buffers were purchased from Fisher Scientific. For comparator data, the same 200 ng injection of K562 were performed on the same system using a 30 minute LC gradient with a ReproSIL C-18 trap and a 15 cm column with 75 μm internal diameter and 1.9 μm bead size. (PepSep). All mass spectrometer settings followed the vendor default method for 1.1 second cycle time DDA, as previously described.(Meier et al., 2018)

### Development of K562 TIMS Folded Spectral Library

The vendor .d file was converted to MGF with MSConvert using the default PASEF MGF settings. The resulting MGF file was processed with the TIMS Folding program placing the 1/k0 value for each peptide at 1/k0 + 100 m/z with an arbitrarily determined intensity of 1,000 counts. Both the original and output files were processed with Proteome Discoverer 2.4 using the IMP-PD community nodes using MSAmanda 2.0 and Percolator for false discovery rate estimation and filtering.(Orsburn, 2021) To accommodate the mass accuracy of the TIMSTOF instrument all files were processed with a precursor tolerance of 30ppm and an MS/MS tolerance of 0.05 Da against the UniProt SwissProt human database downloaded in April of 2021. Oxidation of methionine and protein N-terminal acetylation were used as potential variable modifications and the carbamidomethylation of cysteine was set as a static modification. Target decoy FDR filtering was used at both the peptide and protein level according to manufacturer’s default “Basic Consensus Workflow” template. The resulting .pdresult file from both the original and folded output file were separately imported into Skyline 20.2.0.243 using the default cutoffs to generate a .blib spectral library.(MacLean et al., 2010; Schilling et al., 2012) EncyclopeDIA 1.2.2 (Searle et al., 2018)was used to convert the .blib library to .dlib and to the historic NIST format .MSP. The .MSP libraries were imported into Proteome Discoverer 2.4 or 2.5 our used in the freely available MSPepSearch 2017 command line program.(Lam et al., 2007, 2007, 2008)

### Processing of K562 Data

For comparator data, the folded and nonfolded versions of each MGF were processed identically in Proteome Discoverer 2.5 using MSAmanda 2.0 as described above. The two files were also processed with MSPepSearch separately using both the original spectral library and the library containing folded ion mobility data. Additionally, both MGF files were processed with multiple search engines using SearchGUI and PeptideShaker.(Barsnes and Vaudel, 2018; Vaudel et al., 2015)

#### Data Availability

All files, spectral libraries and processed .msf files are available via ProteomeXchange as dataset PXD029447. Files may be obtained directly from the MASSIVE repository at the following link: ftp://massive.ucsd.edu/MSV000088290/.

## Results and Discussion

### Software compatibility with folded MGF files

To test whether differences were observed between MGF files with and without TIMSFolding values, files MGF files were processed under identical conditions with multiple historic data processing software tools. A summary of some results are shown in Table 1.

**Table 1.** A summary of search tools and the results obtained when using a TIMTOF MGF with and without the TIMS value folded into the MS/MS spectra.

|  | SEARCH ENGINE |  | MGF | FOLDED |
| --- | --- | --- | --- | --- |
| **K562 2 HOUR 200**<br>**μL/MIN** | MSAmanda 2.0 (Dorfer et al., 2014) | Proteins | 5980 | 5956 |
|  |  | Peptides | 39563 | 39495 |
|  |  | PSMs | 48282 | 48209 |
| **K562 30 MIN 500 μL/MIN** | MSPepSearch | Proteins | 3284 | 3283 |
|  |  | Peptides | 13392 | 13388 |
|  |  | PSMs | 17083 | 17075 |
|  | Comet (Eng et al., 2013) | Proteins | 2726 | 2721 |
|  |  | PSMs | 14873 | 14931 |

While there were subtle differences in the number of PSM, peptide, and protein group identifications, in no case was an alteration exceeding 0.8% observed in numbers of identifications. To directly compare the identifications observed, a K562 digest with 30 min LCMS run time was converted to MGF with and without TIMSFolding at with the 1/k0 value added to a fictional peak at 100 m/z with an arbitrarily determined intensity of 1,000 counts.

**Figure 1.**
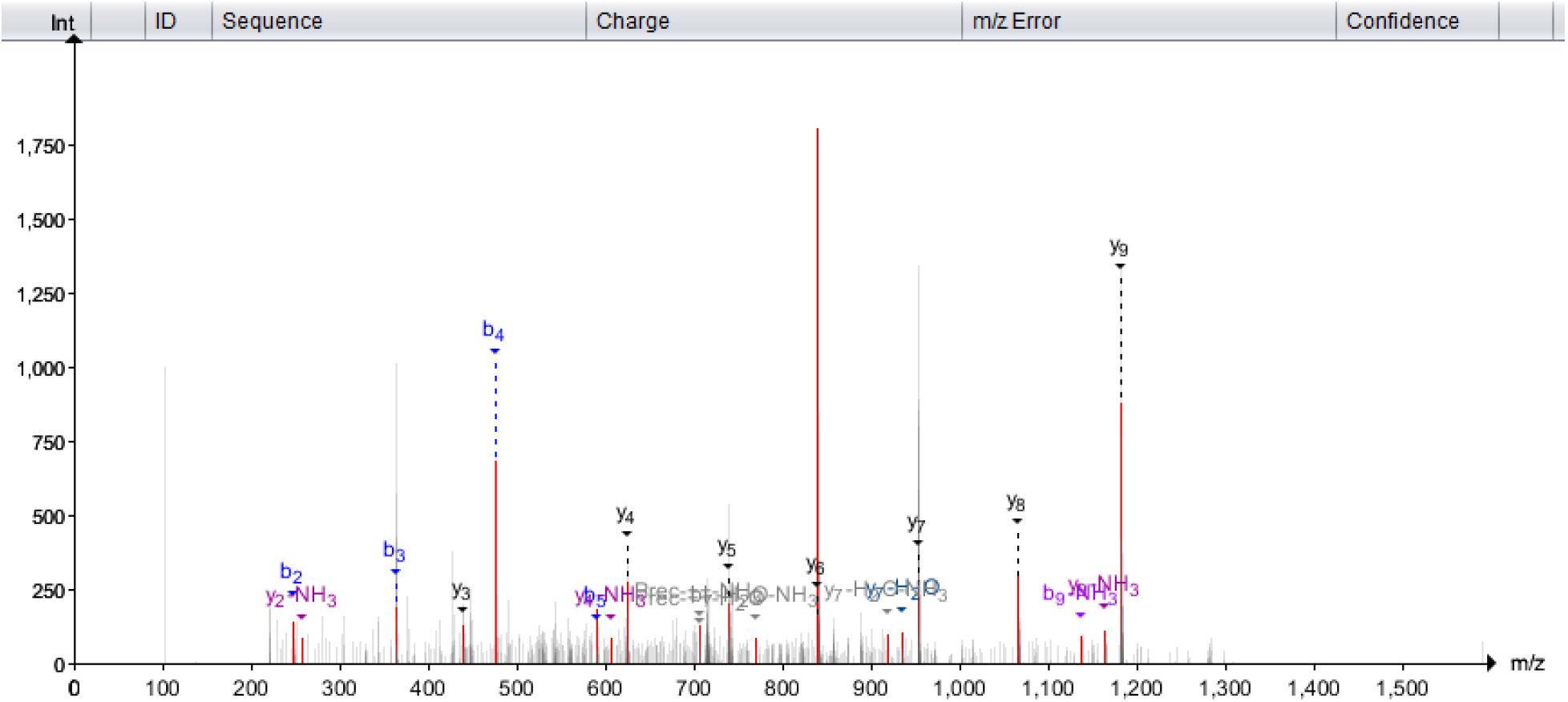
An example peptide spectral match using a TIMSFolded MGF in the multi-search engine software environment of PeptideShaker. The folded 1/k0 value can be observed at approximately 100 m/z in the spectrum.

The original and folded file were processed identically using MSAmanda 2.0 in IMP-Proteome Discoverer 2.4 and using Percolator for FDR estimation and filtering. As shown in Figure 2, the folding process imparted an approximately 0.8% alteration in the peptide and protein identities observed. Protein identifications that differed were largely driven by protein group identifications. For example, P63267 was only identified in the folded MGF search. This protein was identified by a single unique peptide with only three fragment ions identified within the 0.05 Da search tolerance defined.

**Figure 2.**
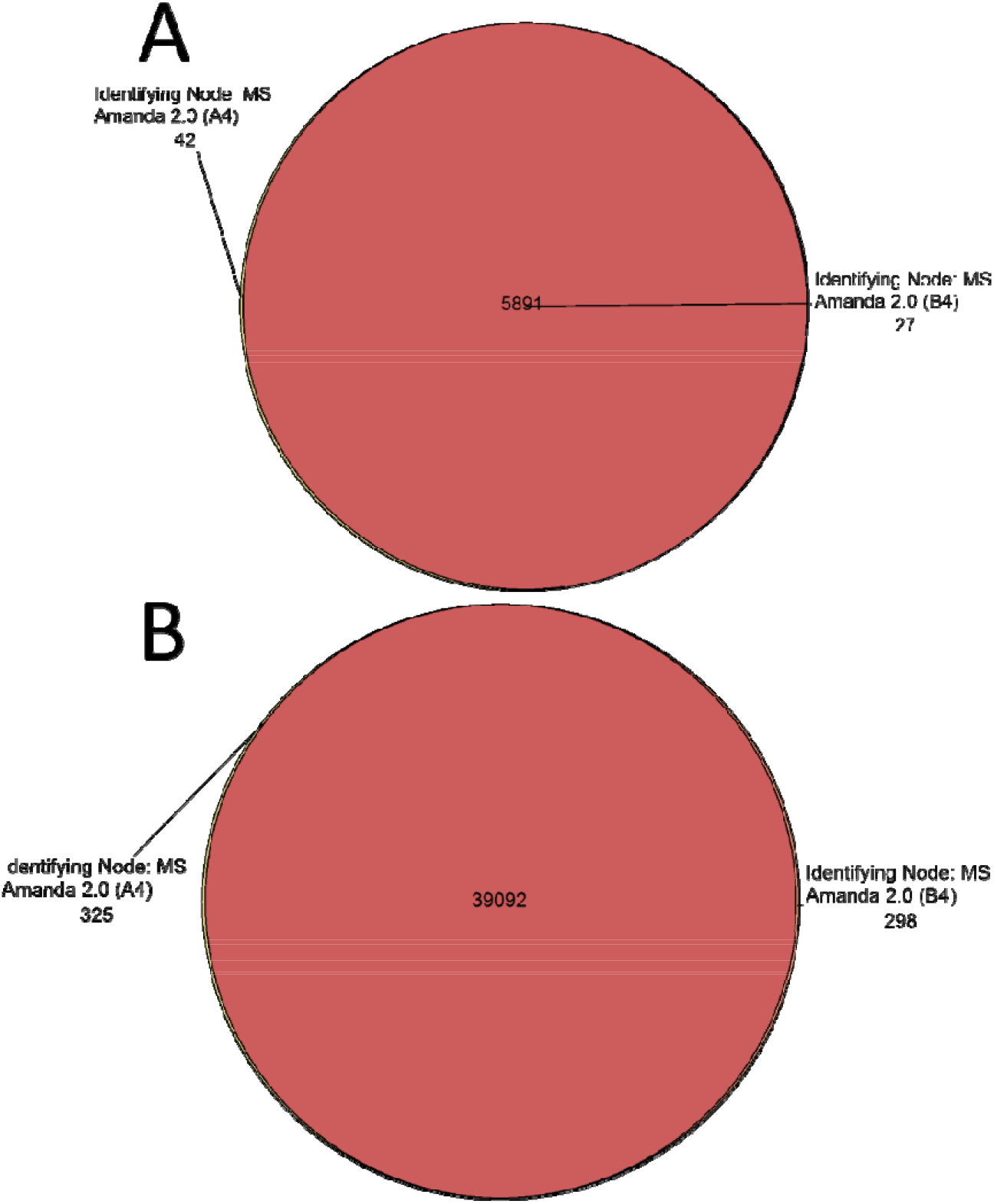
Venn diagrams for (A) the protein group identifications and (B) peptides identified when using the same search parameters and MGF files with (A4) and without (B4) TIMSFolding employed.

### Use of folded ion mobility for an additional confidence metric

As reported by others, the 1/k0 values obtained by the TIMSTOF instruments can be both reliable and predictable for each peptide, as demonstrated by the use of deep learning models.(Meier et al., 2020; Ogata and Ishihama, 2020; Vasilopoulou et al., 2019) As such, the recorded 1/k0 value for each peptide can be used as an additional confidence metric for assigning a peptide identification. Figure 3 is an example peptide spectral match with low fragment sequence confidence and a corresponding 1/k0 measurement that is > 0.073 units off of the library spectra value.

**Figure 3.**
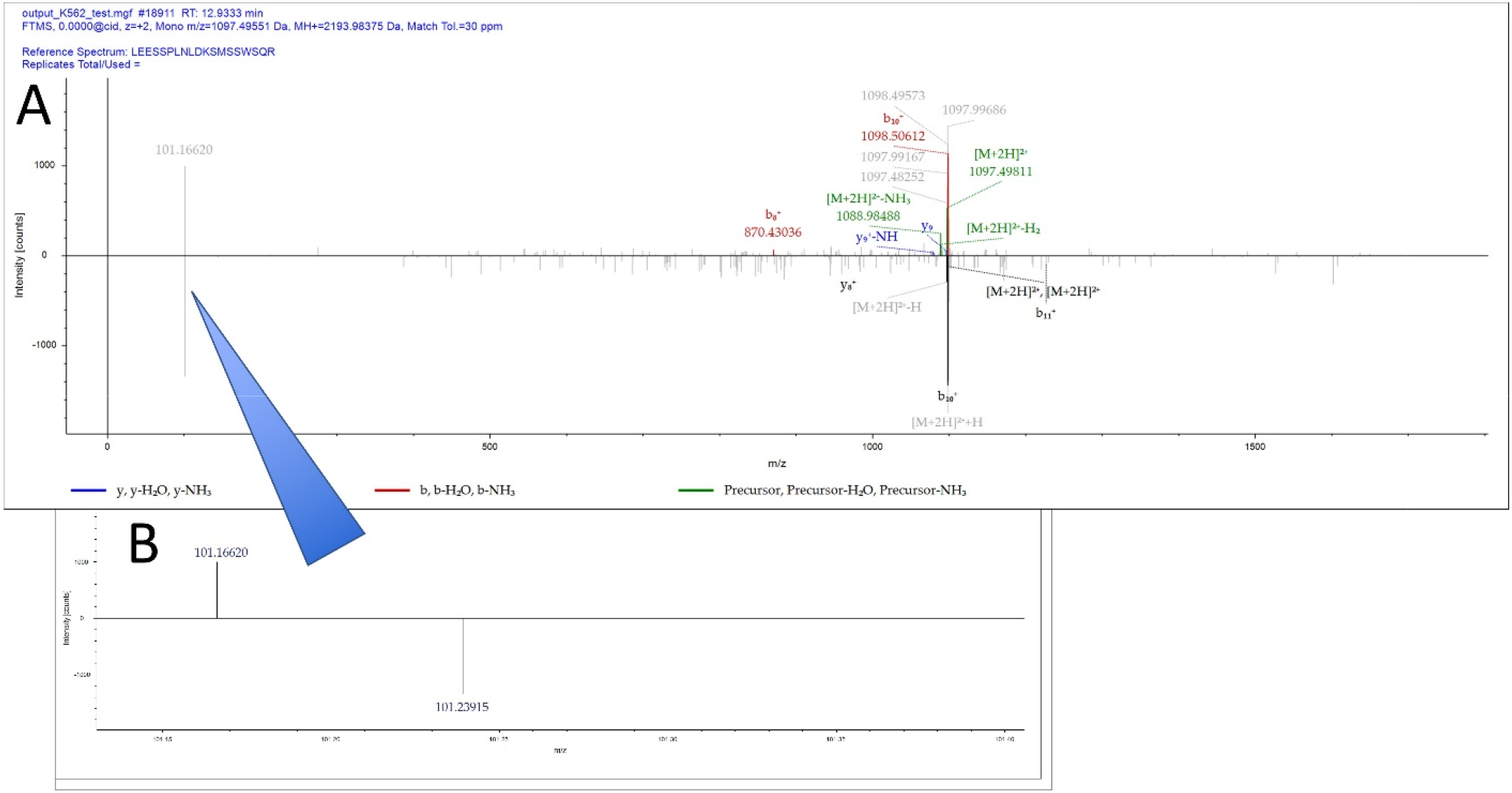
**(A)** An example of a poor match by peptide sequence coverage by MSPepSearch with experimental spectra on top and library spectra on bottom. **(B)** A zoomed in region displaying the TIMSFolding value from the experimental spectra on top and recorded TIMSFolding value on the bottom demonstrating a corresponding mismatch in 1/k0 values.

**Figure 4.**
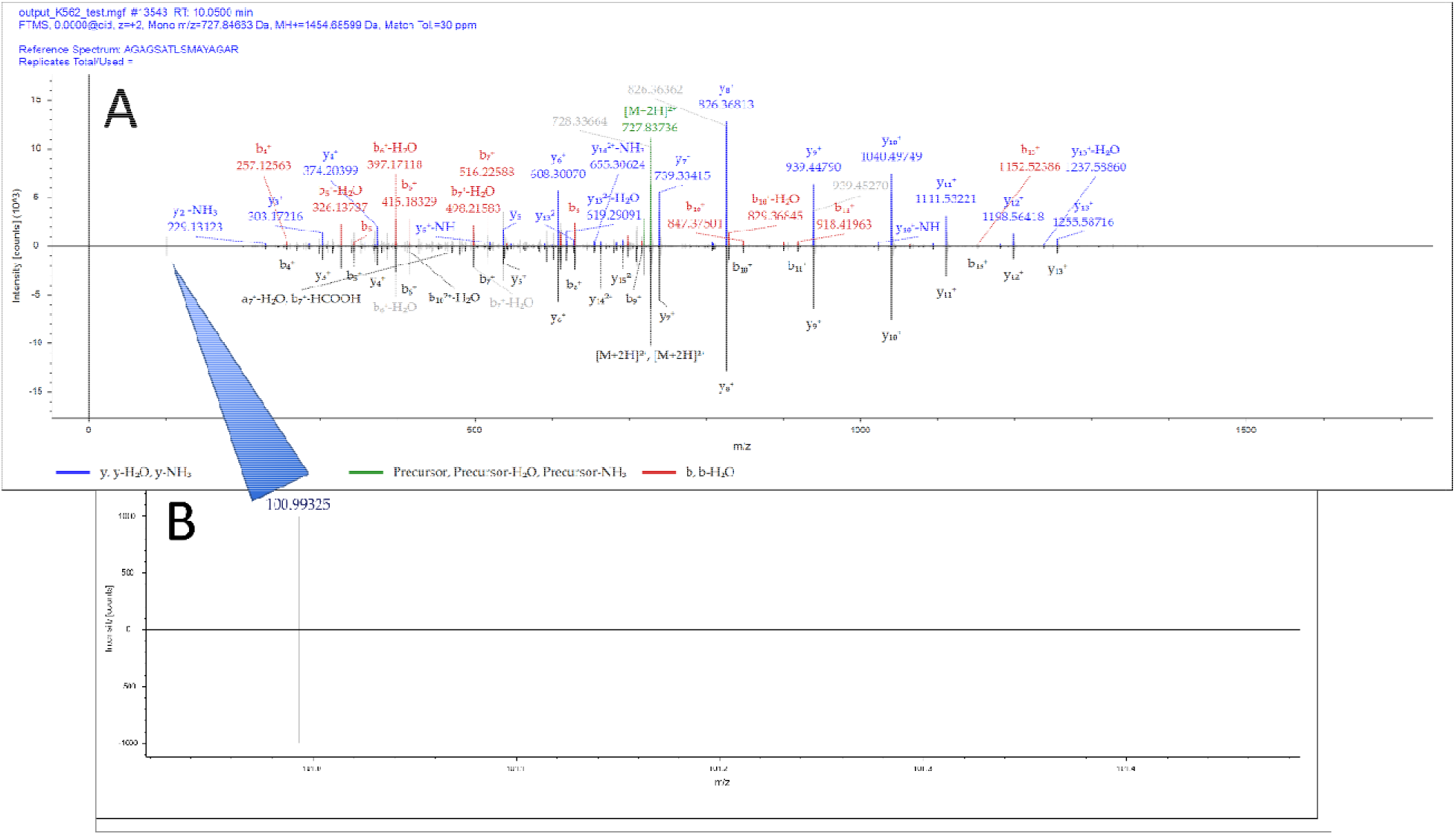
**(A)** An example of a peptide spectral match with 100% sequence coverage with experimental spectra on top and library spectra on bottom. **(B)** The zoomed in region displaying the TIMSFolding values with near perfect match between experimental and library.

## Conclusions

Metcalfe’s law is a generalization of the increasing value of interconnected computer networks argues an exponentially increasing value of a network as nodes are connected.(Friend and Norman, 2013) It is tempting to observe a similar pattern in analytical instrumentation as users develop new applications and resources on each new platform as it emerges. In this context, the rate at which the TIMSTOF technology is continuing to expand in terms of the number of applications presented to date suggests a bright future for this technology as the user base continues to expand. As our lab has adopted TIMSTOF as a central platform, our largest single challenge was a lack of compatibility with both open and commercial data processing tools where we have established investment and expertise. TIMSFolding is the first example of tools in development in our lab to circumvent these challenges and leverage the additional dimension of data this instrument provides compared to our other hardware. Further developments are currently underway, including efforts to automate the creation of TIMSFolded files during data acquisition and to leverage the 1/k0 value as a confidence metric in bulk processed data.

